# Sensory input to cortex encoded on low-dimensional periphery-correlated subspaces

**DOI:** 10.1101/2022.06.15.496327

**Authors:** Andrea K Barreiro, Antonio J Fontenele, Cheng Ly, Prashant Raju, Shree Hari Gautam, Woodrow L Shew

## Abstract

As information about the world is conveyed from the sensory periphery to central neural circuits, it mixes with complex ongoing cortical activity. How do neural populations keep track of sensory signals, separating them from noisy ongoing activity? Here we show that sensory signals are encoded more reliably in certain low-dimensional subspaces. These coding subspaces are defined by correlations between neural activity in primary sensory cortex and upstream sensory brain regions; the most correlated dimensions were best for decoding. We analytically show that these correlation-based coding subspaces improve, reaching optimal limits (without an ideal observer) as noise correlations between cortex and upstream regions are reduced. We show that this principle generalizes across diverse sensory stimuli in the olfactory system and the visual system of awake mice. Our results demonstrate an algorithm the cortex may use to multiplex different functions, processing sensory input in low dimensional subspaces separate from other ongoing functions.

**Significance statement:** Traditionally, primary sensory cortex was thought to have one job – processing sensory signals. As technical advances allow more holistic measurements of the brain and body in action, it has become clear that primary sensory cortex is involved with many other aspects of brain function, not just dealing with sensory input. How can a single neural circuit juggle multiple jobs simultaneously? Here we use numerical, analytical, and experimental methods to demonstrate an algorithm the brain may use to solve this problem by separating different jobs into different subspaces defined by correlations between primary sensory cortex and the brain regions that source the sensory input signals.

## Introduction

Neurons in primary sensory cortices are involved in diverse aspects of brain function; their activity is not limited to encoding sensory signals^1–4^. It is becoming increasingly clear that primary sensory cortex is a multiplex, full of cross talk and multipurpose signals. For example, neuronal activity in primary visual cortex (V1) does not just encode physical features of visual stimuli, but is also related to locomotion^5,6^, whisking and pupil diameter^7^, forepaw manipulations^8^, decision making^8–10^, and learned consequences (rewards) of the visual stimuli^11^. Similarly, neurons in primary olfactory cortex (piriform cortex, PC) go beyond odor coding, exhibiting activity related to spatial navigation^12^, thirst^13^, decision making^10^, and working memory^14^, and can drive distinct behaviors^15^. In general, involvement in these diverse ‘non-sensory’ functions will vary across repeated trials of a sensory stimulus. Thus, it is not surprising that the responses of single cortical neurons to a repeated sensory stimulus vary greatly from trial to trial, often making the stimulus identity impossible to decode accurately with a single neuron. How does the brain reliably keep track of sensory signals when they are mixed into the complex, multipurpose dynamics of the cortex?

Here we propose a population-level solution to this problem. We start from the fact that at the sensory periphery, neuronal activity is purely sensory and not mixed with other functions. As the signal traverses the sensory hierarchy from the periphery to cortex, it becomes increasingly mixed with non-sensory signals due to increasingly recurrent interactions with other brain regions^16,17^. It stands to reason that sensory signals in thalamic nuclei or olfactory bulb could be less noisy (closer to purely sensory) than sensory signals in cortex. Consistent with this, dorsal lateral geniculate nucleus (LGN), which provides input to V1, exhibits response to visual stimuli that has lower dimensionality than V1^18^ and is less affected by locomotion than response in V1^6^. Similarly, LGN firing is modulated more by sensory input and less by behavioral context compared to V1^16^. Likewise, olfactory bulb (OB), which innervates PC, is often less noisy than PC. For example OB has more neurons that are clearly responsive to olfactory stimuli compared with PC^19^. (In Supporting Fig S1, we directly show that the population-level signal-to-noise ratio is greater in OB compared to PC and greater in LGN compared to V1 for the data analyzed further below.) Thus, we hypothesized that certain coding subspaces in cortex – those that share variability with subspaces in upstream sensory regions (thalamus or OB) – may contain sensory signals with less noise.

What do we mean by a coding subspace? Considering a population, rather than single neurons^20,21^, the single-trial response of N cortical neurons is a vector in an N-dimensional space; the sixth component the vector is the response of the sixth neuron, and so on. The responses to many repeated trials of two different stimuli can be represented as two clouds of points in N-dimensional cortical space, one point for each trial, one cloud for each stimulus type. The spread of each cloud of points reflects the trial-to-trial variability (the non-sensory signals discussed above) and overlap of the two clouds makes decoding the stimuli difficult. However, if the response variability due to non-sensory noise lies along different directions than the variability due to switching the sensory signal, then decoding can be greatly enhanced by projecting the N-dimensional response onto a coding subspace, i.e. a lower dimensional subspace that excludes some noise. Our hypothesis here is that such coding subspaces can be found by considering signal correlations and noise correlations between the cortical population and upstream populations.

Recent studies have adopted conceptually related approaches demonstrating that high dimensional neural circuits may manage multiple operations by performing them in different subspaces. For instance, neurons in mouse auditory cortex ‘rotate’ sensory representations from a sensory subspace to a memory subspace over time^22^. Neurons in rat posterior parietal cortex use different subspaces to represent decision and movement^23^. Monkeys making a choice about motion and color of a visual stimulus, exhibited neurons in PFC that used three different subspaces to encode color, motion, and choice^24^. Similarly, working memory and movement planning are separated into different subspaces within a population of lPFC neurons in monkeys^25^. In anesthetized monkeys, signals are transmitted between V1 and V2 in a ‘communication subspace’^26^. The outputs of mouse cerebellar neurons were shown to represent quiescent and active behavioral states in orthogonal subspaces^27^. Computational models together with human brain imaging suggests that orthogonal subspaces are used to represent different task variables in an image classification task^28^. Our work here extends these ideas, establishing sensory subspaces in cortex and in the sensory brain regions that provide input to cortex and new ways to find these subspaces.

Projecting high-dimensional activity into a lower dimensional coding subspace is a type of dimensionality reduction. More generally, dimensionality reduction has long been recognized and used to improve decoding of sensory signals with supervised pattern classification techniques like linear discriminant analysis (LDA) ^29,30^. However, LDA and similar techniques are biologically implausible coding strategies because these algorithms require information about which stimuli caused which response; the brain is not privy to stimulus identities. Without information about stimulus identities, the brain would have to blindly try many (infinite) possible low dimensional projections to find the optimal one, which is implausible. Here we identify a more biologically plausible strategy; we show that low-dimensional optimal coding subspaces can be found without knowledge of stimuli identities by considering correlations between cortex and upstream brain areas that provide input to cortex. Using canonical correlation analysis (CCA, see refs ^21,31^ for an introduction), we define subspaces in cortex and subspaces in LGN or OB in which responses to stimuli are most correlated across the brain regions. We show that these cross-population correlated subspaces can effectively separate signal from noise, often approaching theoretical limits of optimal decoders (like LDA). We developed an analytical approach to better understand these coding subspaces and successfully predicted improved coding subspaces among neurons with low cross-population noise correlations. We first present the theory and then test its predictions using the spike data.

## Results

A central idea underpinning our theory is that the brain can improve decoding of sensory input by projecting neural activity onto a subspace which excludes some of the “noise” that compromises decoding. We hypothesized that we could identify such decoding subspaces based on inter-regional correlations between cortex (V1 or PC) and upstream extracortical regions (LGN or OB). To demonstrate how this might work, we first present a simple, instructive case based on simulated data: two neurons in cortex (Fig 1C) and two neurons in the upstream region (Fig 1B). In this simulated example, the responses are drawn from a multivariate Gaussian distribution (Methods) with parameters chosen such that the two cortical neurons have strong noise correlations and a small difference in mean response for the two stimuli. The two extracortical neurons have noisy overlapping responses to the two stimuli. (In this example, there are no cross-population noise correlations, which is important for our approach, as we discuss further below.) All four of these neurons are rather poor decoders at the single neuron level, but decoding improves substantially when projected onto a particular subspace (green lines in Fig 1B,C). The optimal subspace can easily be found using LDA (dashed line in Fig 1C is the LDA classification boundary), but LDA requires knowledge of the stimulus identities, which is “cheating”; the brain does not have access to stimulus identities before they are decoded. The optimal subspace can also be found, without knowledge of stimulus identity, by performing CCA, which is the key advance presented in this paper.

**Figure 1.**
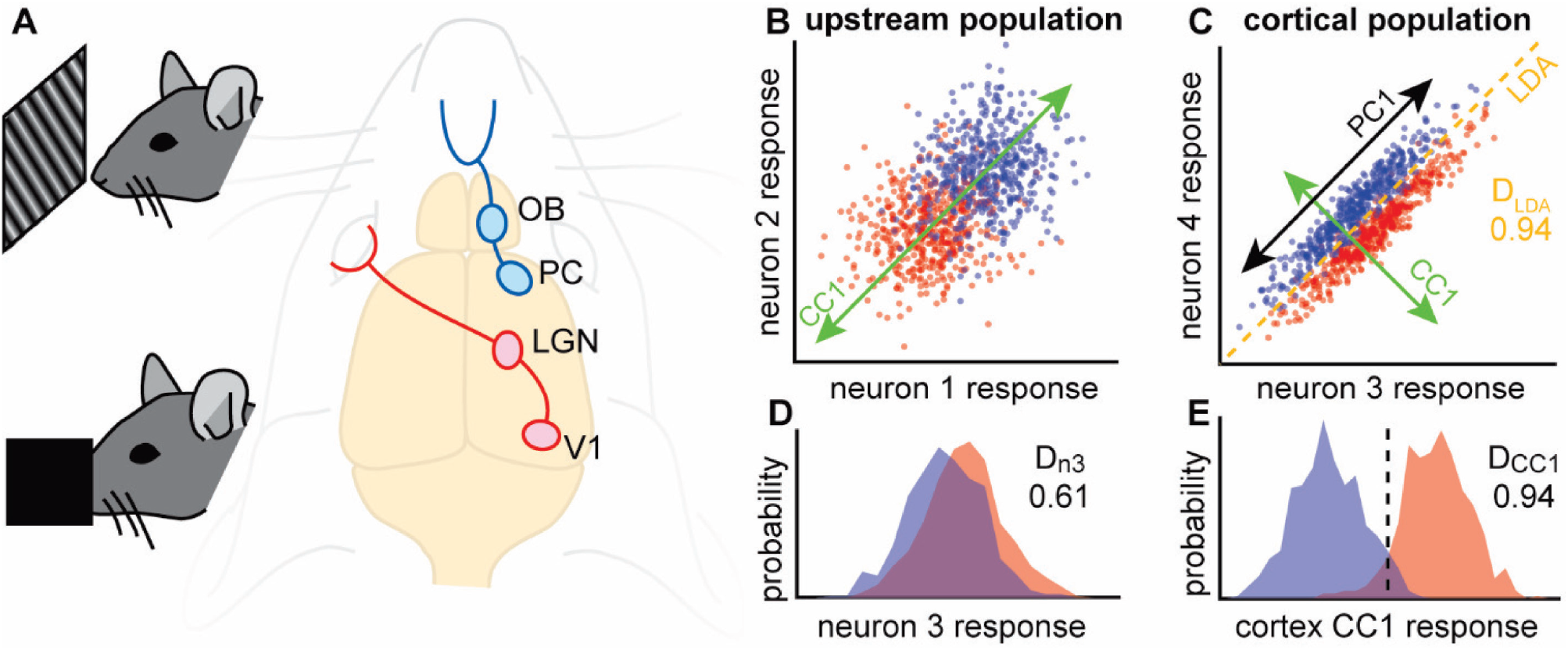
Using inter-regional correlations to find coding subspace. **A)** We hypothesize that correlations between primary sensory cortices (PC or V1) and upstream populations (OB or LGN) can be used (via CCA) to identify coding subspaces with reduced noise. We test this hypothesis in awake mice using simultaneous recordings from V1 and LGN during visual stimulation (top) or PC and OB during olfactory stimulation (bottom). **B)** Each point represents the response of neurons 1 and 2 to one of two stimuli (red and blue). **C)** Responses of neurons 3 and 4 in the cortical population. The optimal LDA decoder achieves 94% accuracy. CCA applied to the two populations identifies a linear subspace (CC1, green) for each population. Projection of each population’s response onto its CC1 results in maximized correlation across populations. PCA finds the linear subspace (PC1, black) with maximal variability; in this case the variability is due to noise. **D)** Response distributions for the two stimulus types overlap substantially for neuron 3, resulting in suboptimal decoding accuracy (61%). **E)** When projected onto CC1, the response distributions better separate the two stimulus types, achieving optimal decoding accuracy (94%, same as LDA). Dashed line indicates the optimal threshold used for calculating the decoding accuracy.

Before proceeding, we briefly introduce CCA for unfamiliar readers (see also refs ^21,31^), comparing and contrasting with the more commonly used principal component analysis (PCA). Similar to PCA, CCA generates a set of basis vectors based on the covariance matrix of multivariate data; these are the canonical components for CCA and the principal components for PCA. In our context, PCA would take a set of spike count responses from a single population of neurons and generate one set of components. In contrast, two sets of spike count responses, one from each of two different populations of neurons, are the inputs to CCA. Likewise, CCA generates two sets of components, one for each population. The first canonical component (CC1) for the first population is related to CC1 for the second population; CCA is defined such that the correlation between the two populations is maximized when they are projected onto their respective CC1s. In contrast, PCA is defined such that projection onto PC1 maximizes variance. In the example in Fig 1C, PC1 (black arrow) is aligned with noise fluctuations in the cortical population, but CC1 (green arrow) is aligned with the direction along which signal varies most (orthogonal to PC1 in this case), thus identifying the optimal decoding subspace (orthogonal to the LDA classification boundary). In general, PCA will not identify the optimal decoding subspace.

Throughout this paper we report various types of decoding accuracy including: D_LDA_, D_n3_, D_CC1_, and optimal D. In all of these cases, we are working with a one-dimensional response to two stimuli; it is one-dimensional because it is projected onto a line or, in the case of D_n3_, simply because it is the response of one single neuron. We define decoding accuracy as the fraction of correctly predicted stimuli using the best possible threshold to separate the responses. For example, in Fig 1E, the best possible threshold is marked with the dashed line.

Is the example in Fig 1 indicative of a more general principle? Does CC1 always reveal the optimal decoding subspace? To answer these questions, we next performed a more extensive analytical and numerical study of this 2×2 case considering a wide variety of correlations among the 4 neurons and signal-to-noise scenarios. We assumed that each neuron had responses to two different stimuli that were drawn from multivariate gaussian distributions. We further assumed that the responses of the 4 neurons are governed by 15 parameters: 8 mean responses (2 stimuli x 4 means), 4 variances, 2 within-population covariance, and 1 cross-population covariance (Fig 2A). Variances and covariances were assumed to be the same for the two stimuli. By centering on the mean response for one stimulus type, we reduce this to 11 parameters, without loss of generality. We considered 50000 different configurations of these 11 parameters, drawn randomly (Methods). We note that here we used analytical methods to compute the CC directions, optimal D, and other quantities; these results depend only on the 11 parameters discussed above and are not limited by finite numbers of samples (Methods).

**Figure 2.**
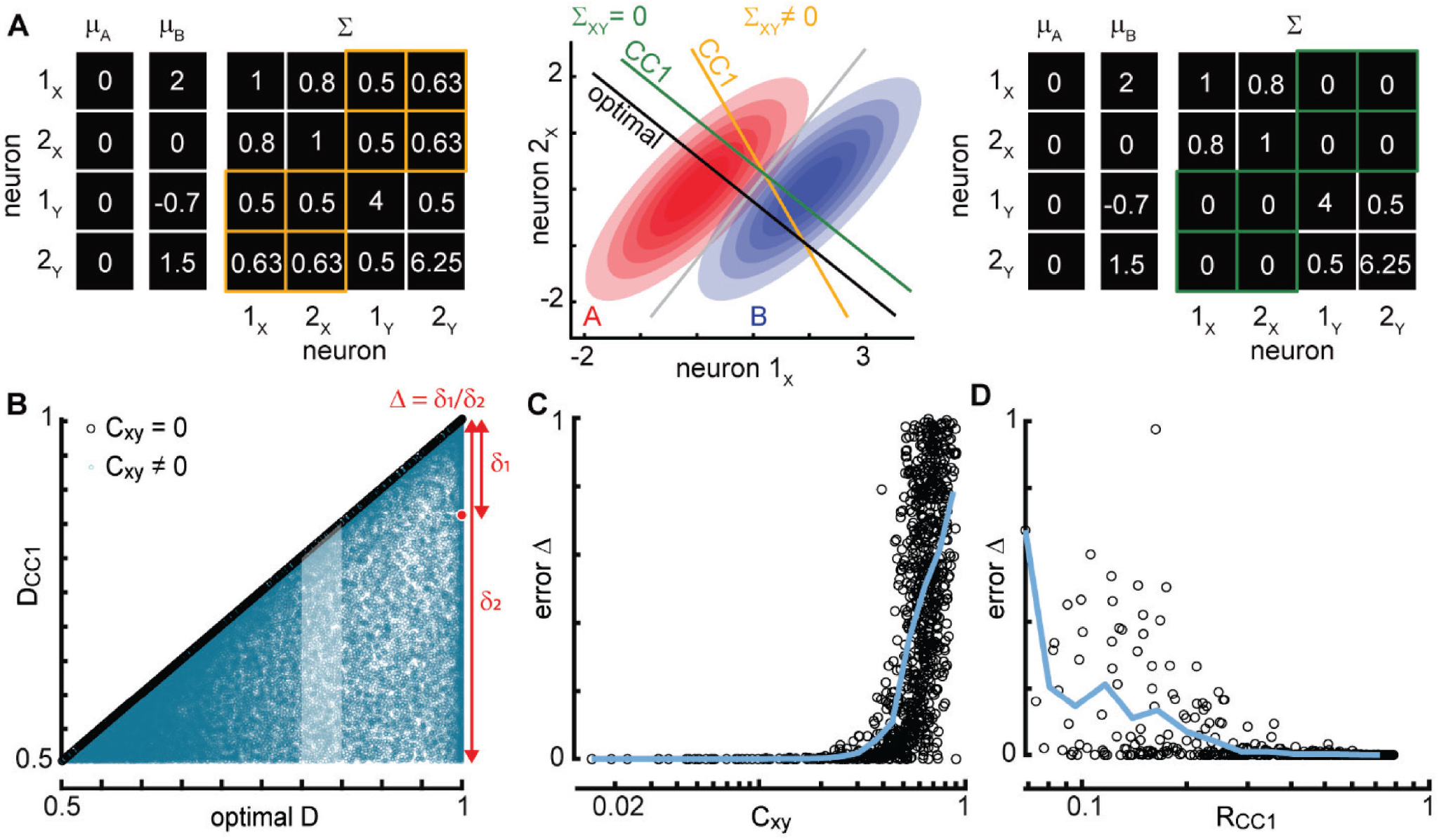
Cross-population noise correlations and CC1 correlation determine accuracy of coding subspace. **A)** Example parameters for population x (neurons 1_x_ and 2_x_) and population y (neurons 1_y_ and 2_y_) for two stimulus types (A and B). When stimulus-independent cross-population covariance between populations x and y are zero (green), the first component CC1 is the same as optimal decoding dimension. Non-zero cross-population covariance results in deviations from the optimal decoding dimension. **B)** Each point represents one of 50000 randomly chosen parameter sets for the 2×2 population. D_CC1_ is perfect (same as optimal D, Δ = 0) if the cross-population noise correlations are zero (black points). If cross-population noise correlations are non-zero, then CC1 decoding is often suboptimal (blue points, Δ > 0). The highlighted set of blue points is studied further in panels C and D. **C)** Decoding error (Δ) is the normalized distance from optimal D (see red diagram in panel B). D_CC1_ is near optimal (low Δ) for C_xy_ below about 0.4. The points shown in panel C exclude cases with very small R_CC1_ (Methods). **D)** Achieving low decoding error Δ also requires sufficiently large R_CC1_. Shown results exclude cases with large C_xy_ (Mehtods). Blue lines in panels C and D – moving average of points.

We found that CC1 is not, in general, well-aligned with the optimal decoding subspace, resulting in decoding D_CC1_ that is often suboptimal (Fig 2B, blue). Nonetheless, for many of the 50000 random populations, D_CC1_ was very close to optimal. Next, we asked what factors determine whether the subspace defined by CC1 is near optimal or not? We found that the most important factor was correlated noise shared across the two populations. When we set these cross-population noise correlations to zero, keeping all the other parameters fixed, the CC1 direction was exactly optimal in all cases (Fig 2B, black and Supporting Information (Mathematical Results)). Thus, our theory predicts that if there were no stimulus-independent shared variability between, say, LGN and V1 neurons, then CCA would be a perfect algorithm for visual decoding, even without knowing the stimulus labels. But, of course, it is very unlikely that LGN and V1 have zero noise correlations. Therefore, we next asked whether near-optimal CC1 decoding might be achieved even with small, non-zero noise correlations? Or, does sensitivity to noise correlations render the algorithm useless in real neural systems? To answer these questions, we examined how D_CC1_ depends on cross-population noise correlations C_xy_. We found that non-zero, but small C_xy_ - below about 0.4 - resulted in a CC1 that was very close to the optimal subspace (Fig 2C). Thus, the algorithm is robust to some degree of noise correlations and could be useful for real sensory systems.

One further prediction from the theory comes from considering the correlation coefficient R_CC1_ of the responses across the two populations after projection onto the two CC1 directions. R_CC1_ is the quantity that is maximized by the CCA algorithm. Note that R_CC1_ includes contributions from both noise and signal-related correlations. We found that if R_CC1_ is too low, then CC1 decoding deteriorates (Fig 2D). In this case, the two populations are nearly independent with no shared information about noise or signal. Thus, a more complete prediction from our theory is as follows. If there exist neurons in primary sensory cortex and the upstream brain regions that have sufficiently low C_xy_ and sufficiently high R_CC1_, then the sensory system could identify low-noise coding subspaces using CCA. Next, we set out to test this prediction in real neurons recorded in awake mice.

Before proceeding, let us examine what it could mean for the experiments to *not* confirm the theory. Why might our predicted CCA decoding subspaces not exist in reality? The most likely reason, as already discussed, is that cross-population noise correlations (C_xy_) could be too large. However, the theory could also fail, because the theory assumed (rather unrealistically) that the responses are Gaussian-distributed and that the neural covariance stricture is the same for both stimuli (Methods). In the experiments, spike count responses are nearly Poisson-distributed and many neurons will certainly have different variance for different stimuli. Finally, it is possible that the limited number of trials in experiments will preclude an effective test of the theory. This would not contradict the theory, but like LDA and other decoding schemes, our CCA decoding scheme when applied to real data with small numbers of trials can, in principle, provide overestimates of decoding accuracy. We will address each of these possible pitfalls below.

To test our predictions, we required simultaneous recordings from at least four neurons, two in primary sensory cortex and two more in an upstream brain region that provides input to cortex. However, considering that many neurons may have substantial noise correlations across populations, our theory predicts that many neurons will be unsuitable for CCA decoding. Therefore, we sought out recordings with far more than 2 neurons in each region. We analyzed two data sets – one in the visual system (V1 and LGN), generated by the Allen Institute^16^, and another in the olfactory system (PC and OB) generated by Bolding and Franks^19,32^. In the Bolding-Franks data (n=8 mice), the number of recorded neurons was 27±6 (mean±SD) in OB and 48±11 in PC. We analyzed responses to six different odorants (fixed concentration) with 15 trials each. In the Allen Institute data (n=9 mice), the number of recorded neurons was 61±27 in LGN and 245±48 in V1. We analyzed responses to static gratings with 6 different orientations and 5 different spatial frequencies with 42 trials each. For both the olfactory and visual data, we defined response as the spike count in the 250 ms period following stimulus onset.

We begin with an example population from the visual system (2 LGN neurons + 2 V1 neurons, Fig 3A) and another example from the olfactory system (2 OB neurons + 2 PC neurons, Fig 3D). For both these examples, projecting response onto CC1 results in optimal decoding for the cortex population. Here optimal decoding is determined with a brute force algorithm; we try all possible lines (with π/200 angular resolution), project responses onto each line, and pick the one with the highest decoding accuracy (Methods). However, considering the same cortical population and choosing a different pair of neurons from LGN (Fig 3B) or a different pair from OB (Fig 3E) can result in a CC1 direction that is far from optimal. Going beyond these example cases, for each mouse and each pair of stimulus types, we considered 10000 randomly chosen populations, each with 2 neurons in cortex and 2 neurons in the upstream region. For each such 2×2 population, we computed the decoding accuracy D_CC1_ for responses projected onto CC1 and compared this to optimal decoding (Fig 3C,F). D_CC1_ was close to optimal for some of these populations, but more typically was far from optimal. Our theory suggests that the populations that deviate from optimal may have large cross-population noise correlations and/or low R_CC1_ (Fig 2). Before testing this prediction, we first verified that our measured values of D_CC1_ are statistically robust (i.e. not artificially high due to overfitting a finite numbers of trials) using a 10-fold cross validation method (Supporting Fig S2).

**Figure 3.**
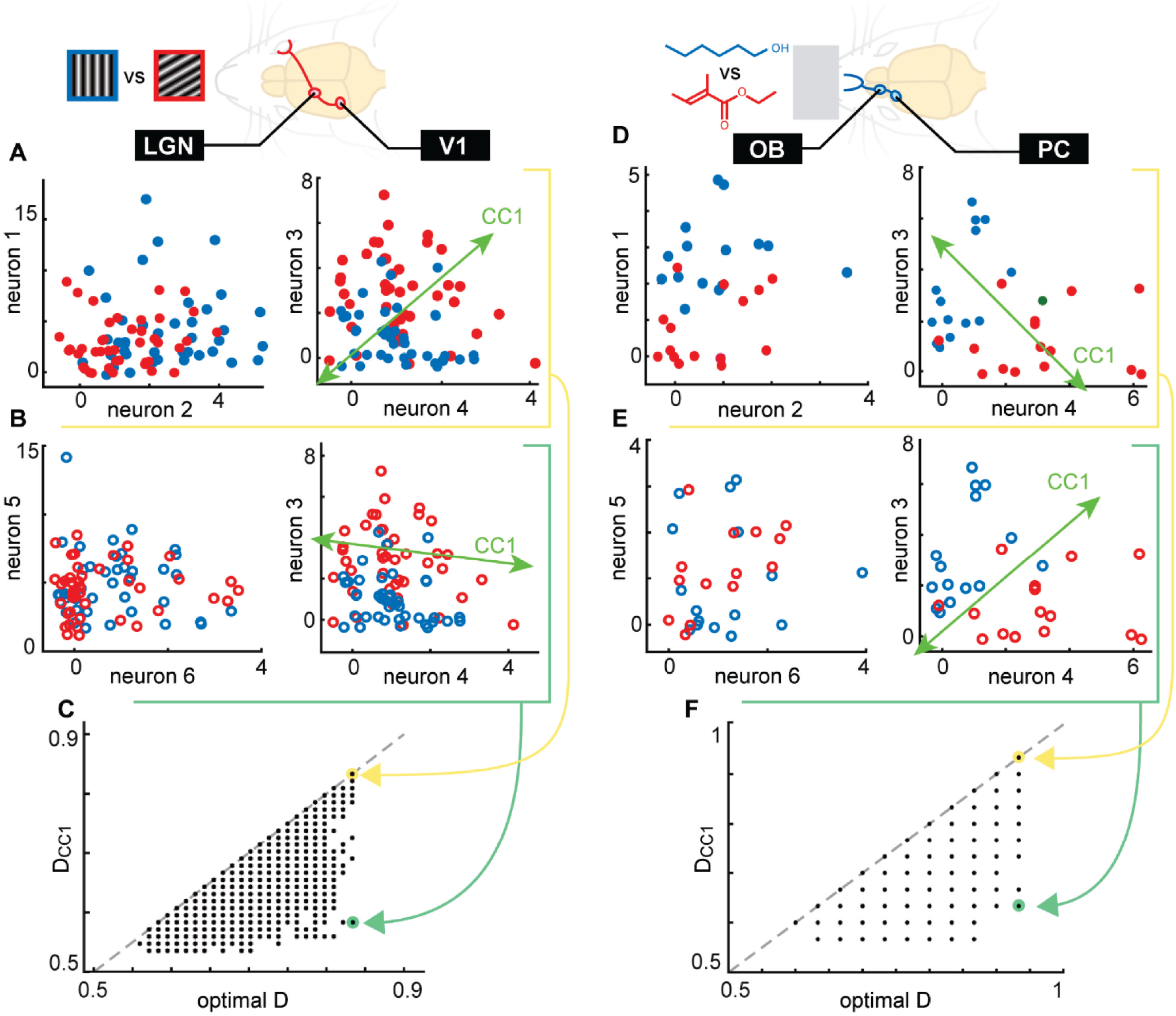
Dual-region neural populations achieve optimal CCA decoding of visual and olfactory stimuli. **A)** Spike count responses from 2 example neurons in LGN (left) and 2 in V1 (right), recorded from an awake mouse viewing two static gratings (90 and 30 degree orientations, 42 trials of each). Projection onto CC1 results in excellent (optimal) decoding in this example. **B)** Same as panel A, except with a different pair of LGN neurons. For this LGN population, the CC1 subspace in cortex becomes suboptimal for decoding. **C)** Summary of CC1 decoding accuracy versus optimal decoding for 10000 randomly chosen 2×2 populations. Green and yellow arrows indicate the examples shown in A,B. **D-F)** Same as panels A-C, but for a different awake mouse breathing two odorants (1-hexanol and ethyl tiglate, 15 trials of each). Note that the discrete values of D are determined by the number of stimulus trials.

To test whether our observed differences between D_CC1_ and optimal decoding were due to high C_XY_ or low R_CC1_, we performed an additional analysis of the populations with high optimal decoding. Our rationale was that these cases can have the largest range of different possible deviations from optimal, and, thus, would be good candidates for studying the source of such deviations. For each mouse and each stimulus pair, we selected the 50 populations (out of the 10000 randomly chosen populations) with the greatest optimal decoding for the two cortical neurons. For each of these, we asked how D_CC1_ for the cortical population varied due to changing the upstream population. We tried 200 randomly chosen upstream populations for each cortical population. In this way, we kept optimal decoding accuracy fixed, while varying D_CC1_, C_xy_, and R_CC1_. In agreement with our theory (Fig 2C), we found that the amount that D_CC1_ deviates from optimal decoding increased as C_xy_ increased for both the visual system (Fig 4A,E) and olfactory system (Fig 4C,G). For this result, we excluded cases with low R_CC1_ (Methods), because our theory predicts that very low R_CC1_ precludes high quality CC1 decoding. As a more direct test of the predicted dependence on R_CC1_ (Fig 2D), we showed that CC1 decoding deviates further from optimal for as R_CC1_ decreases (Fig 4B,D,F,H). Here we excluded cases with high C_xy_ (Methods), because high C_xy_ precludes high quality CC1 decoding. Thus, we conclude that, for many different types of sensory stimulation in many mice, there are multiple neural populations that are consistent with our theory.

**Figure 4.**
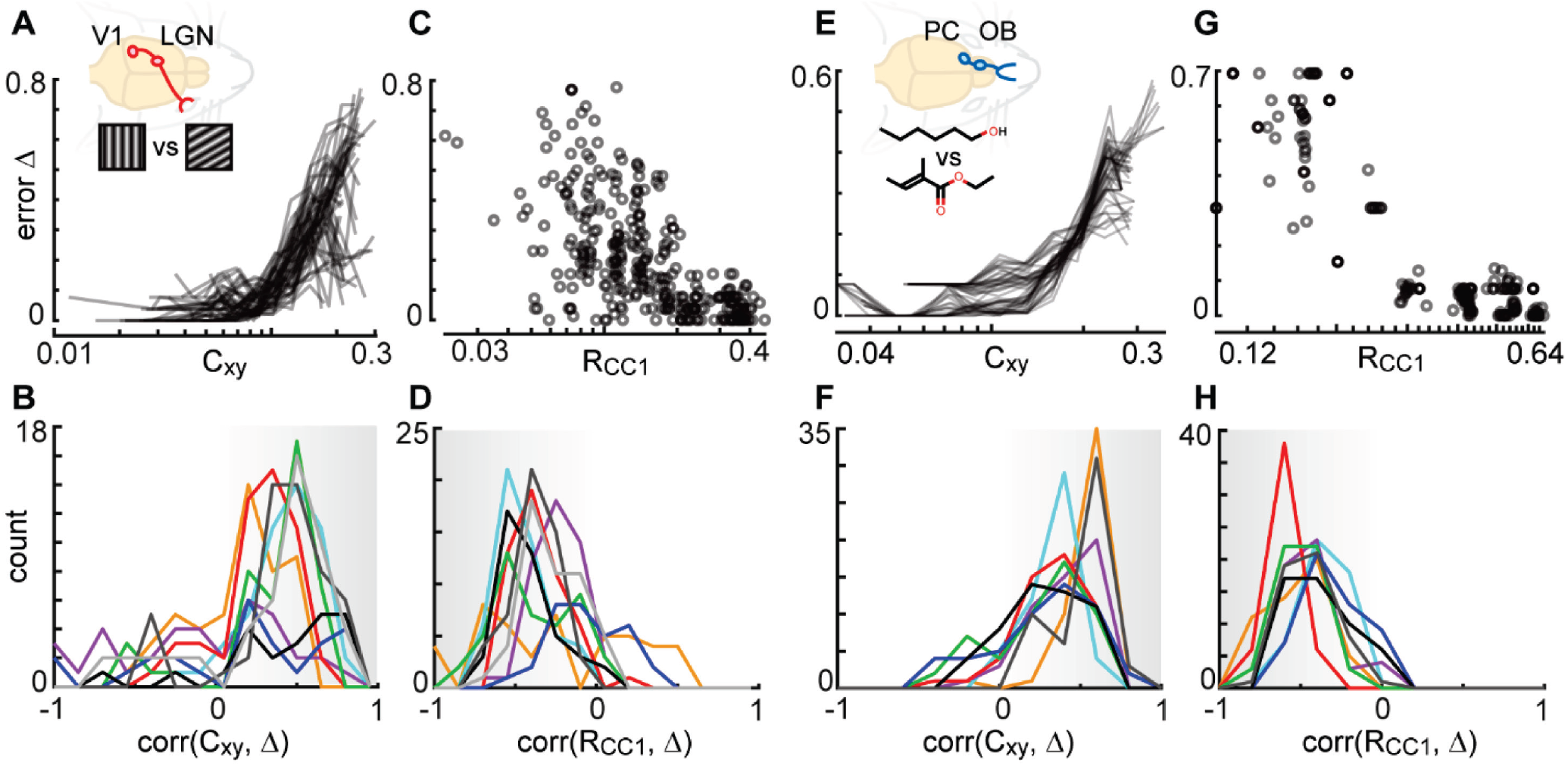
Explaining deviations from optimal CC1 decoding of visual and olfactory stimuli. **A)** As predicted, the difference (error Δ) between optimal decoding and D_CC1_ was often strongly correlated with cross-population noise correlation C_xy_. Each line represents an average over 200 different pairs of LGN neurons and one pair of V1 neurons (excluding those with small R_CC1_, see Methods). **B)** Each distribution summarizes the correlations between error Δ and C_xy_ for 200 populations in one mouse and one pair of stimuli (the color code for specific mouse/stimulus is shown in Supporting Fig S1). Notice that the vast majority of these correlations are positive, like the examples in panel A. **C)** As predicted, error Δ is negatively correlated with R_CC1_. Each point represents one 2×2 population (excluding cases with large C_xy_, see Mehtods). **D)** Each distribution summarizes the anticorrelations between error Δ and R_CC1_ for one mouse and stimulus pair (same cases as panel B). **E-H)** Same as panels A-D, but for OB+PC populations in different mice with olfactory stimulation.

## Discussion

We have described an algorithm – CC1 decoding – that may be used by neural circuits in primary sensory cortex to improve accuracy of sensory decoding, in an “unsupervised” way, i.e. without “ideal observer” knowledge of stimulus labels. The algorithm achieves this decoding improvement by projecting high dimensional population responses onto a one-dimensional subspace with reduced noise (the first canonical component CC1). The low-noise CC1 subspace is readily found by canonical correlation analysis (CCA), provided that noise correlations between cortex and upstream brain regions are not too large and the signals are not too weak. We demonstrated that many real neurons measured in awake mice, in visual and olfactory systems (LGN and V1, OB and PC), meet the conditions required for CC1 decoding.

How might real neurons implement the CC1 decoding algorithm we describe here? Such an implementation requires three operations. First, the CC1 subspace must be identified. Second, neural activity must be projected onto CC1. Third, the projected activity must be thresholded to “decide” which type of stimulus was present. The latter two operations are quite naturally performed by neurons. Neurons sum up their inputs, weighted by synapse strength, which performs projections onto subspaces. For instance, in the cartoon in Fig 5, neurons 3 and 4 together generate a two-dimensional input signal to neuron 5, but when added up and weighted by w3 and w4, their input becomes one-dimensional; it is projected onto a line determined by the relative weights of w3 and w4. Moreover, if neuron 5 also receives input from the upstream population (neurons 1 and 2 in Fig 5), then its input could sum up two projections, one onto CC1 for the upstream population, the other onto CC1 for the cortical population. Recalling that these two projections are maximally correlated, their sum would be an excellent one-dimensional signal for decoding the stimuli. After this projection, the spiking mechanism of neurons naturally thresholds the one-dimensional input. How might neurons identify CC1? In other words, how might they tune the synaptic weights so that they do the specific projections onto CC1? A line of theoretical research has addressed this possibility directly^33–35^ culminating recently^34^ in a model with biologically plausible synaptic plasticity mechanisms. According to this theory, a network of reciprocally connected excitatory and inhibitory neurons that receive input like that received by neuron 5 in Fig 5 can perform precisely the sum of two CC1 projections illustrated in Fig 5. This operation can be performed “on-the-fly”; it does not require a “memory” of previous observations to be stored for later use. The synaptic plasticity rules are based solely on information available locally to each synapse – information about the pre- and post- synaptic neurons’ activity. Moreover, the plasticity rules are based on non-Hebbian mechanisms involving Ca^2+^ plateau potentials^36^ which are consistent with evidence from cortex. Taken together with our work here, we conclude that CC1 decoding not only can reach optimal limits of decoding but is also plausibly performed by cortical circuits.

**Figure 5.**
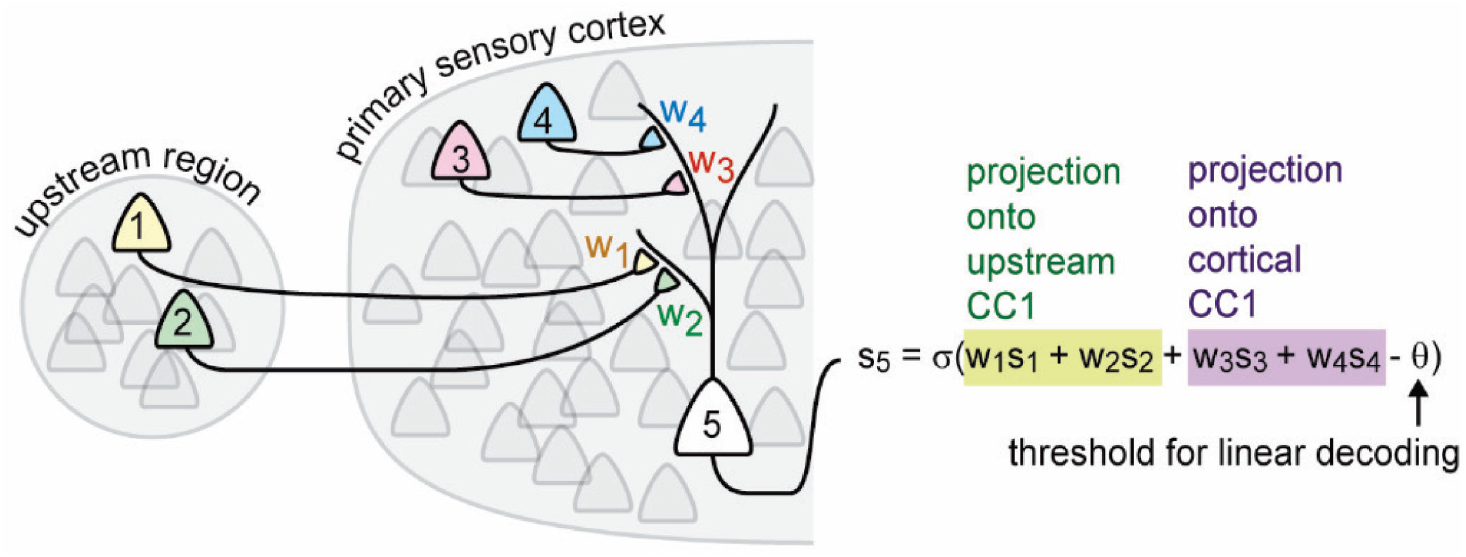
Biophysical implementation of CC1 decoding. Synaptically- weighted summation of inputs naturally performs projections of high-dimensional input onto 1-dimensional subspace. The spike threshold Θ naturally imposes a “decision boundary” for classifying stimulus type of input. Synaptic plasticity mechanisms can tune the synaptic weights so that the projections are aligned with CC1 (Lipshutz et al. 2021).

Here we focused on sensory coding subspaces, but our findings suggest a more general principle for multiplexing many functions within the same neural circuit. Any two brain regions that cooperate to execute a particular function are likely to exhibit some correlated activity. But this shared signal is likely mixed in with other activity (‘noise’) that is involved in other ongoing functions. Our results suggest that the common, correlated activity between the two regions can define a CC1 subspace which effectively separates the function of interest from other ongoing functions, thus allowing the same circuits to execute many functions simultaneously.

## Materials and Methods

### Experiments

The olfactory data set was recorded and first reported in (Bolding & Franks, 2018). Their methods were approved by Duke University Institutional Animal Care and Use Committee. Their methods for olfactory stimulation, head-fixation, respiration monitoring, electrophysiology, and spike-sorting were also described in detail previously^37^. Here studied the following recordings: 170608, 170609, 170613, 170615, 170618, 170619, 170621, 170622.

The visual data set was first reported in (Siegle et al, 2021) recorded by the Allen Institute for Brain Science. The visual stimulation, head-fixation, electrophysiology, and spike-sorting are described and the data is available for public download here: https://allensdk.readthedocs.io/en/latest/visual_coding_neuropixels.html Animal use protocols were approved by the Allen Institute’s Institutional Animal Care and Use Committee. Here we analyzed experiments with the following session IDs: 754312389, 715093703, 750749662, 754829445, 757970808, 759883607, 761418226, 763673393, 799864342.

### CCA with Matlab

To perform CCA on experimental data, we used the Matlab function ‘canoncorr’, e.g. [A,B,∼,U,V,∼]=canoncorr(Region1Data,Region2Data). Here, each set of responses (e.g. Region1Data and Region2Data) is a T x N matrix (T trials, N neurons). The ith trial from Region1Data projected onto the jth CC is stored as U(i,j), and the vector that defines the direction of the jth CC in Region1 is stored as the jth column of A. Note that for the analytical study (Figs. 1 and 2), CCA was not done with Matlab; the CC directions were computed through the associated eigenvalue problem (see below and Supporting Information).

### Optimal decoding

For two-dimensional Poisson-distributed spike count responses, like the experimental data we studied here, typical supervised machine learning algorithms (e.g. LDA and support vector machines) often do not find the optimal decoding accuracy. Thus, we made a brute-force algorithm that projects the responses onto a line and evaluates decoding accuracy (as described in next subsection), trying many possible angles of projection between 0 and π rad (in steps of π/200 rad). Optimal decoding accuracy is that of the projection with the maximum decoding accuracy of all these projections.

### Decoding accuracy

The decoding accuracy of various linear projections are reported in the main text (optimal D, D_CC1_, D_n3_). After responses were projected down to one dimension, the decoding accuracy was determined by trying every possible threshold between the minimum and maximum projected response. The fraction of correct classifications for the best threshold was reported as decoding accuracy. For the analytical study of 2×2 populations a different approach (independent of number of trials) was taken (see below and Supporting Material.)

### Predictions about C_xy_ and R_CC1_

In Fig 2C,D, we showed that D_CC1_ approaches optimal decoding for small C_xy_ and large R_CC1_. To better separate the effects of these two quantities, we excluded cases with low R_CC1_ in Fig 2C and excluded cases with high C_xy_ for Fig 2D. These cutoffs were R_CC1_ > 0.75 and C_xy_ < 0.1. We did a similar cutoff for the results in Fig 4, showing that D_CC1_ deviated from optimal for large C_xy_ or small R_CC1_. For Fig 4, the cutoffs were set as follows: for LGN-V1 data, R_CC1_ > 0.3 and C_xy_ < 0.06; for OB-PC data, R_CC1_ > 0.4 and C_xy_ < 0.1.

### Analytical 2×2 study

In Figs 1 and 2, we consider the case of two populations (X and Y) of simulated neurons whose responses to two stimuli, A and B, are correlated both within and across populations. We assume the responses (r_X_ and r_Y_) to each stimulus can be described by a multivariate Gaussian,

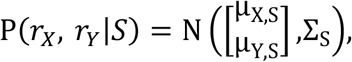

where S = {*A, B*}, μ_*X,S*_ ∈ *R*^*m*^, μ_*Y,S*_ ∈ *R*^*n*^, and Σ_*S*_ is a symmetric, positive-definite matrix of size (*m* + *n*) × (*m* + *n*). Here populations X and Y contain m and n neurons respectively (m = 2 and n = 2 for the simulated results in Figs 1 and 2); for example, Y may be a cortical region and X may represent OB or LGN. Without loss of generality, we simplify notation by shifting the mean responses so that μ_*X, A*_ = 0, μ_*Y, A*_ = 0; thus, we can drop the stimulus subscript on the mean vectors and use μ_*X*_ = μ_*X,B*_, μ_*Y*_ = μ_*Y,B*_. We further assume that the stimulus-conditioned noise covariance matrix is the same for each stimulus: i.e. that Σ_*A*_ = Σ_*B*_ =: Σ. For simplicity, we assume that noise correlations c_xy_ were equal for any pair of cells across the two populations.

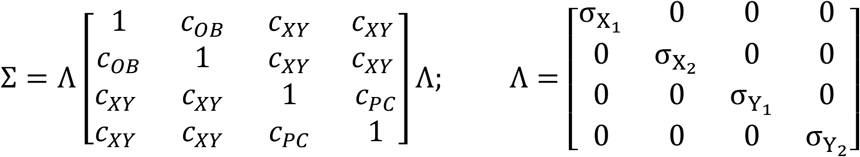

As noted earlier, mean response to stimulus A was 0 and mean response to stimulus B was:

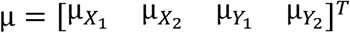

Each of the eleven parameters 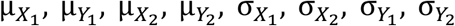, *c*_*OB*_, *c*_*PC*_, *c*_XY_ was chosen randomly from the following distributions:

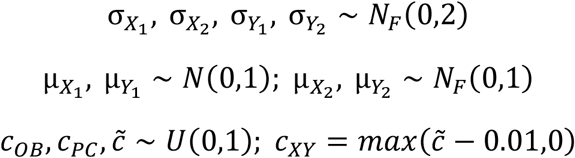

Here *N*_*F*_ is the folded normal distribution (if X ∼ *N*(μ, σ), then |X| ∼ *N*_F_ (μ, σ)), and U is the uniform distribution. Parameters are defined so that – without loss of generality – 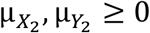, all noise correlation parameters are non-negative, and *c*_*XY*_ has about a 1% chance of being 0. For a single simulation, each parameter was chosen independently from the above distributions; the covariance matrix was then checked for positive-definiteness (equivalently, 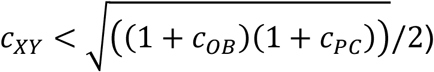: if it failed, a new set of parameters was chosen. This was then repeated 50000 times, allowing a robust and wide-ranging survey of possible signal and noise correlation structures.

We next explored decoding under different assumptions. First, we sought to determine how well the stimulus can be decoded from responses within each population. In this simplified setting (responses are Gaussian, and the noise covariance is stimulus-independent), the optimal decoder is linear and can be determined by a simple analytical formula (see Supporting Information, Methods). That is, we decode the stimulus by projecting the population response onto a single vector, and then compare that value with a threshold. Next, we use the principal direction from canonical correlation analysis, or CC1, as a linear decoder. Finally, we artificially remove cross-population noise correlations by setting *c*_*XY*_ = 0, and recompute the canonical correlation analysis with the revised stimulus-unconditioned covariance. We show that in this setting, the most correlated direction CC1 is in fact equal to the optimal projection vector (see Supporting Information, Methods). These three decoding vectors – optimal, CC1, and CC1 with *c*_*XY*_ = 0 – are demonstrated in Fig. 5A as black, yellow and green respectively.

We computed the single-population optimal decoding directions *v*_*X*_, *v*_*Y*_ using Eqn. (S1) and found the decoding accuracy by integrating the resulting one-dimensional Gaussians (see Eqn. (S2)). Similarly, we calculated the CC1 for each population using Eqn. (S4), and found the decoding accuracy using *v*_*X,CC*1_, *v*_*Y,CC*1_ as projection vectors. To compute single-cell decoding accuracy, we integrated under the marginal distributions given by projecting onto the coordinate directions 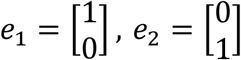.

## Supporting Information for

### Supporting Information Text

#### Supplemental Experimental Results

**Fig. S1.**
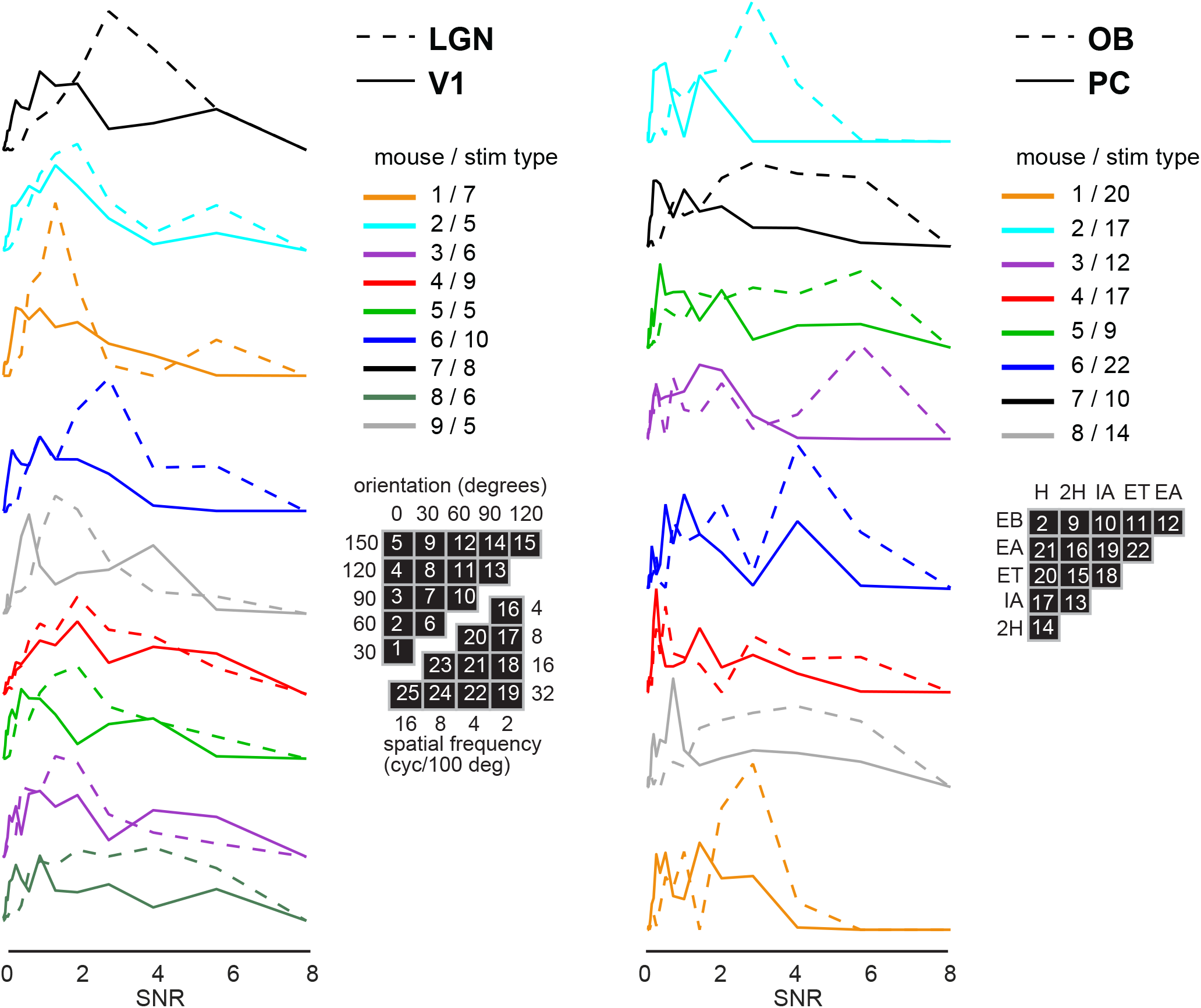
Signal-to-noise ratio is lower in cortex than upstream regions. In the main text, it was suggested that signal-to-noise is lower in primary sensory cortex compared with the upstream regions, closer to the sensory periphery, that provide sensory input to cortex. Here we show quantitatively that this is true for the visual system and olfactory system data that we analyzed. [Left] Each solid line is a distribution of SNR values for the 10000 randomly chosen pairs of V1 neurons analyzed in the main text. Each dashed line is a SNR distribution for the corresponding 10000 pairs of LGN neurons. Here we took the 2×2-population-level SNR to be 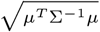, where *μ* is a two-element vector of response differences (response to stimulus type A minus response to stimulus type B) and Σ is the response covariance matrix for the two neurons. Each color represents an example from a different mouse and a different pair of stimuli. The ‘stim type’ number in the legend refers to a specific pair of grating orientations or spatial frequencies as labeled in the black grids. [Right] Same as the left, but based on the olfactory system. Note that for both the visual and the olfactory systems, the solid distributions indicate a lower typical SNR in cortex compared to the upstream regions (dashed). Also note that this set of example mice and stimulus types is the same as those shown in Fig 4B,D,F and H, with the same color code.

**Fig. S2.**
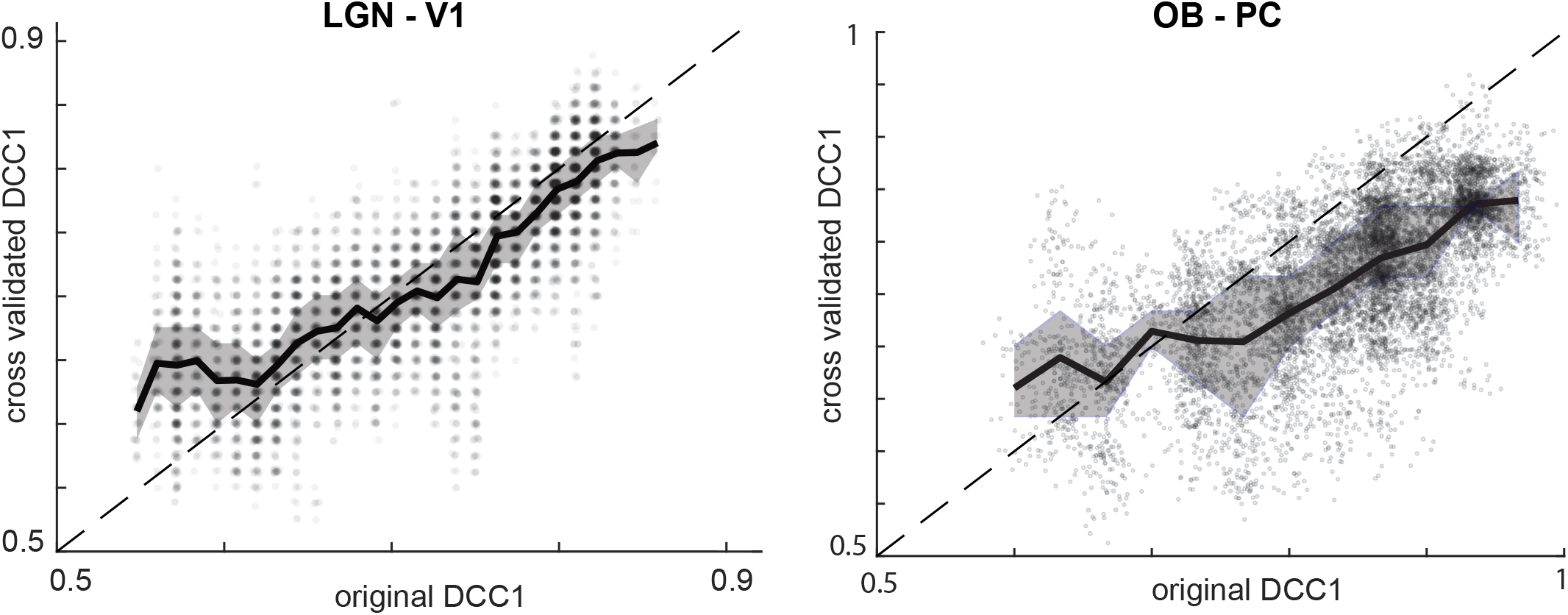
Cross-validation of DCC1. Like many other decoding algorithms, our proposed CC1 decoding can generate artificially high values of decoding accuracy if there are too few stimulus trials or if response dimensionality is too high. In the context of our work, response dimensionality is simply the number of neurons from each population, which was 2 for all the results reported here. Although our theory is not limited to this 2-dimensional case, reliable tests of the theory using a finite number of trials require a low dimensional response. We tried 3×3 populations and found (not shown) that the cross-validated DCC1 became less reliable than shown here. For this reason, we chose to work with 2×2 populations. For our 2×2 populations, we did a 10-fold cross validation to verify that our measured values of DCC1 were reliable. For a given 2×2 population, we used 9/10 of the stimulus trials to calculate CC1 directions and determine the optimal decoding threshold on the CC1-projected responses. Then, we used the same projection and threshold to calculate decoding accuracy for the held out 1/10 of trials. We repeated this for the 10 unique folds of training and hold-out trials and finally averaged the 10 decoding accuracy values across the 10 folds. These averaged decoding accuracies are reported here compared to the original decoding accuracies reported in the main text. Note that the cross validated DCC1 values are strongly correlated with the original values, which means they are reliable. However, it is clear that for the olfactory data (right), which had only 15 trials per stimulus type, original DCC1 is more prone to over estimation compared the cross validated DCC1. The visual system data (left) included 42 trials per stimulus type, which allows a better match between original and cross validated DCC1. The dashed line (slope unity) marks equality between original and cross validated DCC1. Each point represents one of 10000 randomly chosen populations. The dark line and shaded area indicates the median and quartiles of the points, respectively. Points are displayed with 20% opacity, so that density of points is clearer.

### Mathematical Results

#### Introduction

We consider the case of two populations of neurons whose responses to two stimuli, A and B, are correlated both within and across populations. We assume the responses to each stimulus can be described by a multivariate Gaussian, i.e.

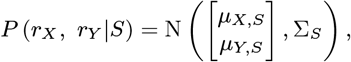

where *S* = {*A,B*}, *μ*_*X,S*_ ∈ ℝ^*m*^, *μ*_*Y,S*_ ∈ ℝ^*n*^, and Σ_*S*_ is a symmetric, positive-definite matrix of size (*m* + *n*) × (*m* + *n*). Here X and Y refer to two populations of neurons which are both responsive to A and B, containing m and n neurons respectively; for example, Y may be a cortical region and X a pre-cortical region which supplies afferent input to Y. Without loss of generality, we simplify notation by shifting the mean responses so that *μ*_*X,A*_ = 0, *μ*_*Y,A*_ = 0; thus, we can drop the stimulus subscript on the mean vectors and use *μ*_*X*_ = *μ*_*X,B*_ and *μ*_*Y*_ = *μ*_*Y,B*_.

We next consider how to decode the stimulus, using only the responses within each population. We assume that the covariance matrix is the same for both stimuli: i.e. Σ_*A*_ = Σ_*B*_ =: Σ. In this case the optimal decision boundary is given by a hyperplane in ℝ^*m*^ or ℝ^*n*^; equivalently, by a one-dimensional projection of the response vector. The decision boundary is given by (for example) *u* ∈ ℝ^*m*^ such that 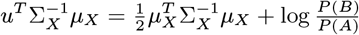, (Here, Σ_*X*_ and Σ_*Y*_ are the marginal covariances in populations X and Y respectively.) Therefore, the projection vector must be the normal vector to this plane; i.e.:

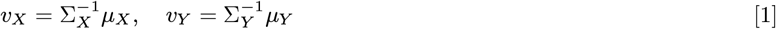

in populations X and Y respectively. Alternatively, observing that *v*^*T*^ *r*_*X*_ |*S* is a one-dimensional Gaussian with

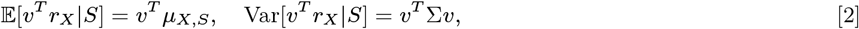

we can derive the same outcome by maximizing the signal-to-noise ratio; i.e. 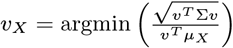. From the perspective of *linear discriminant analysis*, this maximizes between-class (where “class”=stimulus identity) variability while minimizing within-class variability (1).

#### When cross-region noise correlations are absent, CC1 is a perfect decoder

We now compute the projection directions associated with canonical correlation analysis (CCA). Given two sets of zero-mean observations from X and Y, the goal of CCA is to find the linear projections of the observations that are maximally correlated (2). This technique uses the full stimulus-averaged population response; however, we will show that under certain conditions, the maximally correlated direction from CCA coincides with the optimal decoder. Assuming *P* (*A*) = *P* (*B*), the covariance structure within each population is

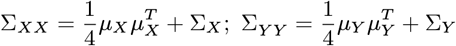

while the stimulus-averaged covariance matrix between populations X and Y is

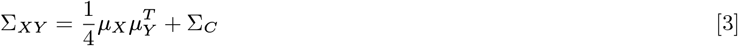

Here Σ_*X*_, Σ_*Y*_, and Σ_*C*_ are the covariances within and across-populations: i.e.

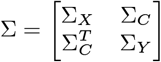

We now seek to find the directions which maximize correlation across the population; that is

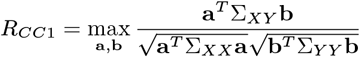

We denote the vectors that achieve this maximum as *v*_*X,CC*1_ and *v*_*Y,CC*1_ respectively; i.e.

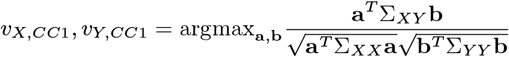

The vectors *v*_*X,CC*1_ and *v*_*Y,CC*1_ can be obtained by finding the principal eigenvectors of *D*_*X*_ and *D*_*Y*_ respectively:

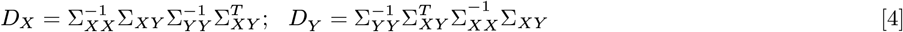

and the corresponding eigenvalue is the correlation (*R*_*CC*1_) squared: that is,

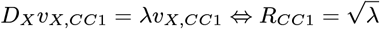

We note that the cross-covariance matrix Σ_*XY*_ has two contributions, one reflecting signal correlations 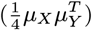 and the other noise correlations (Σ_*C*_). The latter reflects trial-to-trial correlations which are not reflected in the mean response. We will now show that when noise correlations are absent (Σ_*C*_ = 0), the principal CCA direction coincides with the optimal decoding direction. Without loss of generality, we focus on *D*_*X*_ ; parallel statements hold for *D*_*Y*_.

##### Lemma 1

If Σ_*C*_ = 0, then *D*_*X*_ is a rank 1 matrix.

*Proof*. It is well known that the rank of a matrix product is bounded above by the minimum rank of the matrices; i.e. rank(*AB*) ≤ min(rank(*A*), rank(*B*)). From Eq. (3) Σ_*XY*_ is the sum of two matrices, the first of which is rank 1; if Σ_*C*_ = 0, therefore, the sum is rank 1 as well. Therefore any matrix product that includes Σ_*XY*_ has rank at most 1. □

##### Theorem 1

If Σ_*C*_ = 0, then the correlated (non-zero) eigenvector of *D*_*X*_ coincides with the projection direction which is optimal for decoding.

*Proof*. Recall that Eq. (1) shows that 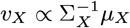. We will show that *v*_*X*_ is also an eigenvector of *D*_*X*_.

Consider the formula for *D*_*X*_:

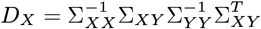

The cross-population correlation matrix Σ_*XY*_ is rank 1 and range(Σ_*XY*_) = Span {*μ*_*X*_}. Therefore, 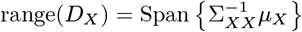.

Next, we write 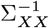 in terms of 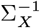. Using the matrix determinant lemma, and noting that

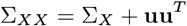

where **u** = *μ*_*X*_*/*2,

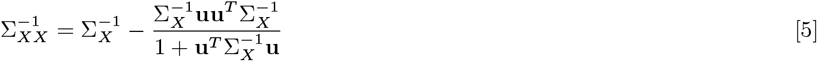

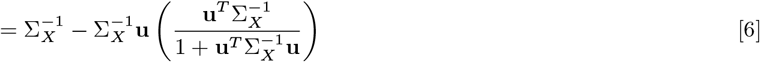

The second term *already* maps into Span 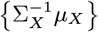, regardless of what vector is multiplied on the right. In conclusion, 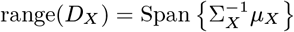; i.e. *D*_*X*_*v*_*X*_ ∝ *v*_*X*_. □

By using Eq. (6) (and the analogous simplification for 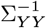), one can confirm that the corresponding eigenvalue is

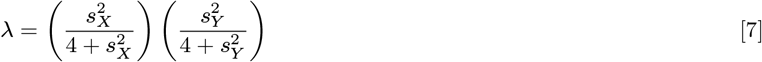

where 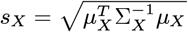 and 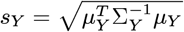 are the signal-to-noise ratios for the *X* and *Y* populations respectively.

##### Theorem 2

If Σ_*C*_ = 0, then any other eigenvector of *D*_*X*_ gives chance-level decoding.

*Proof*. If **v**^*T*^ *μ*_*X*_ = 0, then 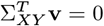 and therefore *D*_*X*_ **v** = 0. Therefore **v** is an eigenvector of *D*_*X*_ with eigenvalue 0. But then

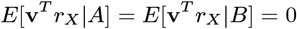

i.e. the stimuli A and B cannot be discriminated. □

